# Microtubule-based perception of mechanical conflicts controls plant organ morphogenesis

**DOI:** 10.1101/2021.09.09.459674

**Authors:** Dorothee Stöckle, Blanca Jazmin Reyes-Hernández, Amaya Vilches Barro, Milica Nenadic, Zsófia Winter, Sophie Marc-Martin, Lotte Bald, Robertas Ursache, Satoshi Fujita, Alexis Maizel, Joop EM Vermeer

## Abstract

Precise coordination between cells and tissues is essential for differential growth in plants. During lateral root formation in Arabidopsis thaliana, the endodermis is actively remodeled to allow outgrowth of the new organ. Here, we show that microtubule arrays facing lateral root founder cells display a higher order compared to arrays on the opposite wall of the same cell, and this asymmetry is required for endodermal remodeling and lateral root initiation. We identify that MICROTUBULE ASSOCIATED PROTEIN 70-5 is necessary for the establishment of this spatially defined microtubule organization and endodermis remodeling, and thus contributes to lateral root morphogenesis. We propose that MAP70-5 and cortical microtubule arrays in the endodermis integrate the mechanical signals generated by lateral root outgrowth, facilitating the channeling of organogenesis.

## MAIN TEXT

Morphogenesis in plants depends on local growth rates and growth directions. Since plant cells are confined and linked together by rigid extracellular cell walls, spatial differences in growth can generate mechanical stresses within tissues, unlike in animal systems. The mechanical tensions caused by cells pulling or pushing on their neighbors are increasingly recognized as instructive signals during development, as well as an important element of the feedback mechanism coupling tissue geometry to gene expression (1-3). The lattice of cortical microtubules (CMTs) plays important roles in translating mechanical signals during morphogenesis (4-6). CMTs align with maximal tensile stress in plant tissues (2, 5) and are required for guiding the cellulose synthase complexes to deposit cellulose microfibrils in the cell wall (7, 8). Therefore, they are important regulators of anisotropic growth at the crossroads of biochemical and mechanical growth control. These conclusions were drawn from the analysis of the epidermal surface of the shoot apical meristem, where cells are under strong tension and not fully differentiated. However, it remains poorly understood how plant cells detect and integrate mechanical signals during de novo morphogenesis within a tissue.

In Arabidopsis thaliana (Arabidopsis), one example of morphogenesis that entails a difference in growth within a tissue is the formation of lateral root primordia (LRP) that initiate deep within the primary root, in the cell file adjacent to the xylem, the xylem pole pericycle (XPP) (9, 10). In response to auxin, lateral root founder cells (LRFCs) swell, their nuclei migrate towards each other, and they divide asymmetrically to form a stage I LRP (11-13). The endodermis that overlies the forming LRP accommodates the radially expanding LRFCs through a change in cell shape and/or volume loss, controlled by AUX/IAA SHORT HYPOCOTYL 2 (SHY2)-mediated endodermal auxin signaling. Interference with this step results in a complete block of LRP formation (14), and the absence of endodermal remodeling. Several lines of evidence support a key role for the remodeling of the endodermis for LR initiation and morphogenesis (14-16), but the nature of the signal perceived by the endodermis upon radial expansion of the LRP remains unknown. Here, we investigate the role of CMTs in the endodermis during LRP formation. We show that the endodermal CMT lattice organization and response are polarized. On the inner side, in contact with the pericycle, the arrays are more-ordered than those on the outer side of the same cell. Specific disruption of CMTs in endodermal cells overlying an LRP result in delayed cellular remodeling and a flattened LRP with atypical cell division patterns. Reorganization of endodermal CMTs depends on both the swelling of the underlying pericycle and a SHY2-dependent auxin response. We identify MICROTUBULE ASSOCIATED PROTEIN 70-5 (MAP70-5) as required for the organization of the endodermal CMT lattice, the remodeling of the endodermis, and the morphogenesis of the LRP. We propose that CMTs and MAP70-5 contribute to the perception of LRP outgrowth by the endodermis and together function as an auxin-regulated integrator of mechanical constraints during organogenesis.

## RESULTS

### Endodermal CMTs reorganize during spatial accommodation

To observe and quantify CMT organization and dynamics in the endodermis during LRP initiation, we generated plants expressing a fluorescent CMT marker, *CASP1pro::mVenus:MBD*. Endodermal cells are highly polarized along the radial axis with two distinct domains, an inner and an outer side, separated by the Casparian strip (17). The inner side of the endodermis is in contact with the radially expanding LRP (**Fig. 1A**). Prior to noticeable endodermal thinning that is required to accommodate LRP development, we observed that CMTs are differentially organized between the inner and outer side of differentiated endodermal cells. CMTs on the inner side form anisotropic arrays oriented along the shoot-root axis, whereas they are more isotropic on the outer side (**Fig. 1B, D**). As the LRP radially expands, the CMT arrays on the inner side reorient and become more isotropic, resembling the CMT organization on the outer side (**Fig. 1C, D**). Together, these results indicate that two spatially defined domains of CMT organization exist in differentiated endodermal cells, and that endodermal CMTs facing the LRP reorganize in response to the radial expansion of the LRFCs during LRP initiation.

**Fig 1.**
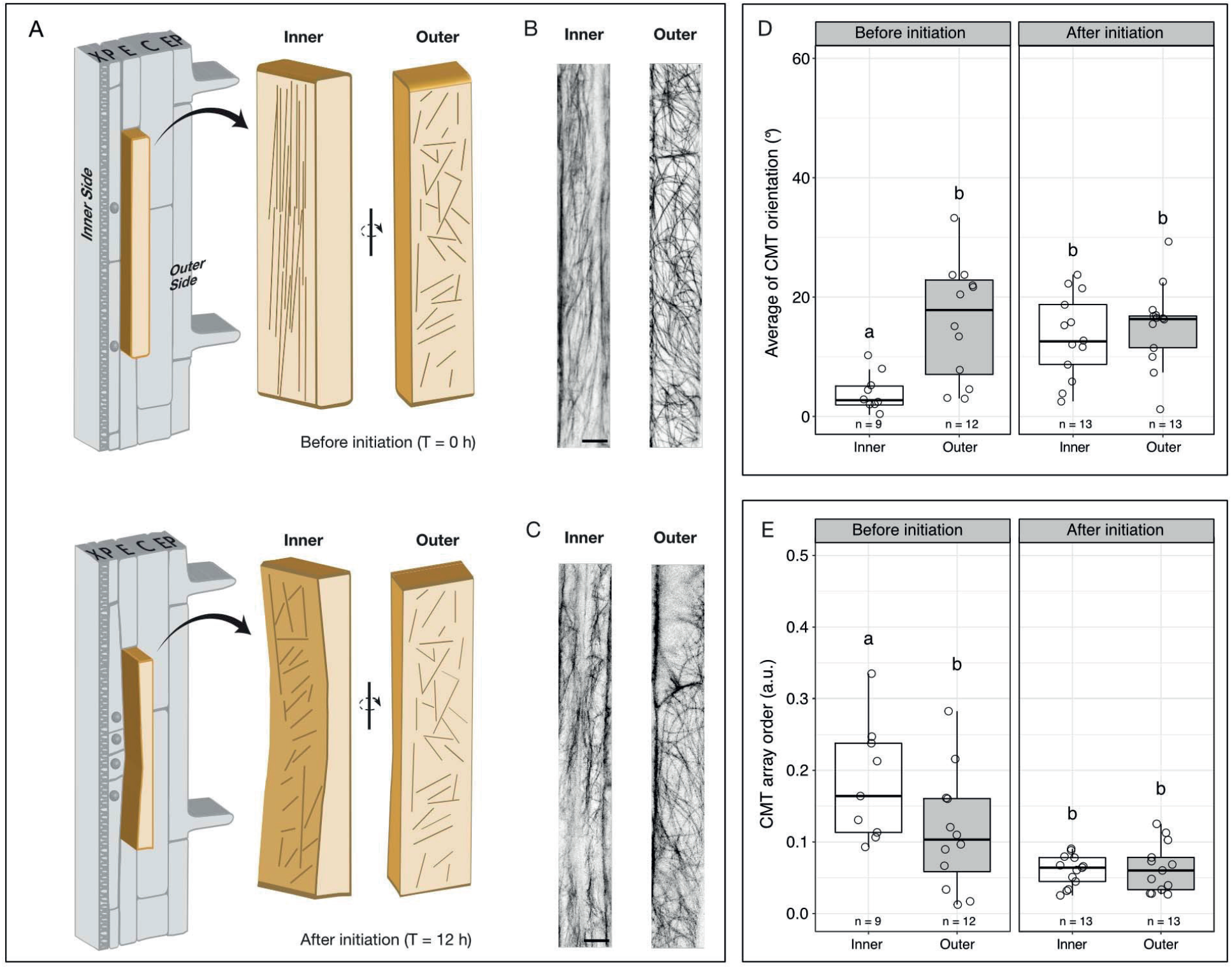
Spatiotemporally-regulated CMT reorganization in overlying endodermal cells during LRP initiation. (**A**) Schematic representation of the remodeling of a differentiated endodermal cell (light brown) overlying a LRFC as it radially expands and divides. The inner and outer sides of the cell remodel differently as the LRP grows (top versus bottom). CMT organization is simplified by lines. (**B, C**) Confocal microscopy images of CASP1pro::mVenus:MBD showing the organization of CMT arrays on the inner and outer side before (t = 0) (**B**) and after (t = 12h) (**C**) gravistimulation. (**D**-**E**). Distribution of CMT orientation before and after LRP initiation. (**D**) Depicts the CMT orientation in degrees with respect to the long axis of the cell, and (**E**) describes the CMT organization on the inner and outer side at indicated time points (arbitrary units 0-1, 0 = no order, 1 = order). X = protoxylem, P = xylem pole pericycle, E = endodermis, C = cortex and Ep = epidermis. Scale bar = 10 µm. Comparison between samples was performed using two-way ANOVA and Tukey’s HSD. Samples with identical letters do not significantly differ (α = 0.05).

### Endodermal thinning and LRP morphogenesis require CMT reorganization

To test whether CMT organization of endodermal cells contributes to the spatial accommodation of the LRP, we took advantage of a truncated version of *PROPYZAMIDE-HYPERSENSITIVE 1* (*PHS1*), *PHS1*Δ*P*, to interfere with the microtubule organization. Inducible, ectopic expression of *PHS1*Δ*P* in cells results in depolymerization of the microtubules (13, 18). We expressed *PHS1*Δ*P* using two complementary endodermis-specific promoters and induction systems: *ELTPpro>>PHS1*Δ*P* and *CASP1pro>>PHS1*Δ*P*. Whereas *ELTPpro* is moderately expressed throughout the whole differentiated endodermis, *CASP1pro* activity peaks early in differentiating endodermal cells during Casparian strip formation (19, 20). This allowed us to analyze the contributions of CMTs during early and later stages of LRP development, and branching in general. We verified that expression of *PHS1*Δ*P* in the endodermis results in the depolymerization of CMTs exclusively in these cells (**Supplemental Fig. S1**). With either expression system, we observed that upon *PHS1*Δ*P* induction, thinning of endodermal cells is delayed and LRP morphology altered (**Fig. 2A-D**). We quantified the impact of perturbing CMTs in the endodermis on LRP development using two different approaches. First, we compared the distribution of LRP developmental stages in the LR development zone (LRDZ), the area encompassing the first visible stage I LRP closest to the root tip and the first emerged LR (stage VIII) (21). *ELTPpro>>PHS1*Δ*P* induced CMT depolymerization resulted in an increased number of LRPs, particularly stage I LRPs (**Fig. 2E, F**). Second, we used gravistimulation-mediated LR formation to quantify differences in the progression of LRP development (22, 23). This revealed that *CASP1pro>>PHS1*Δ*P*-induced CMT depolymerization results in a delay of LRP development (**Fig. 2G, Supplemental Fig. S2**). 36 h after induction of LRP formation by gravistimulation, LRPs are still at stages II/III, while mock treated samples display LRPs at stages VI/VII (**Fig. 2H**). Together, these results show that depolymerization of CMTs in the endodermis prevents its normal remodeling, alters the cell division pattern in the LRP, and delays emergence.

**Fig 2.**
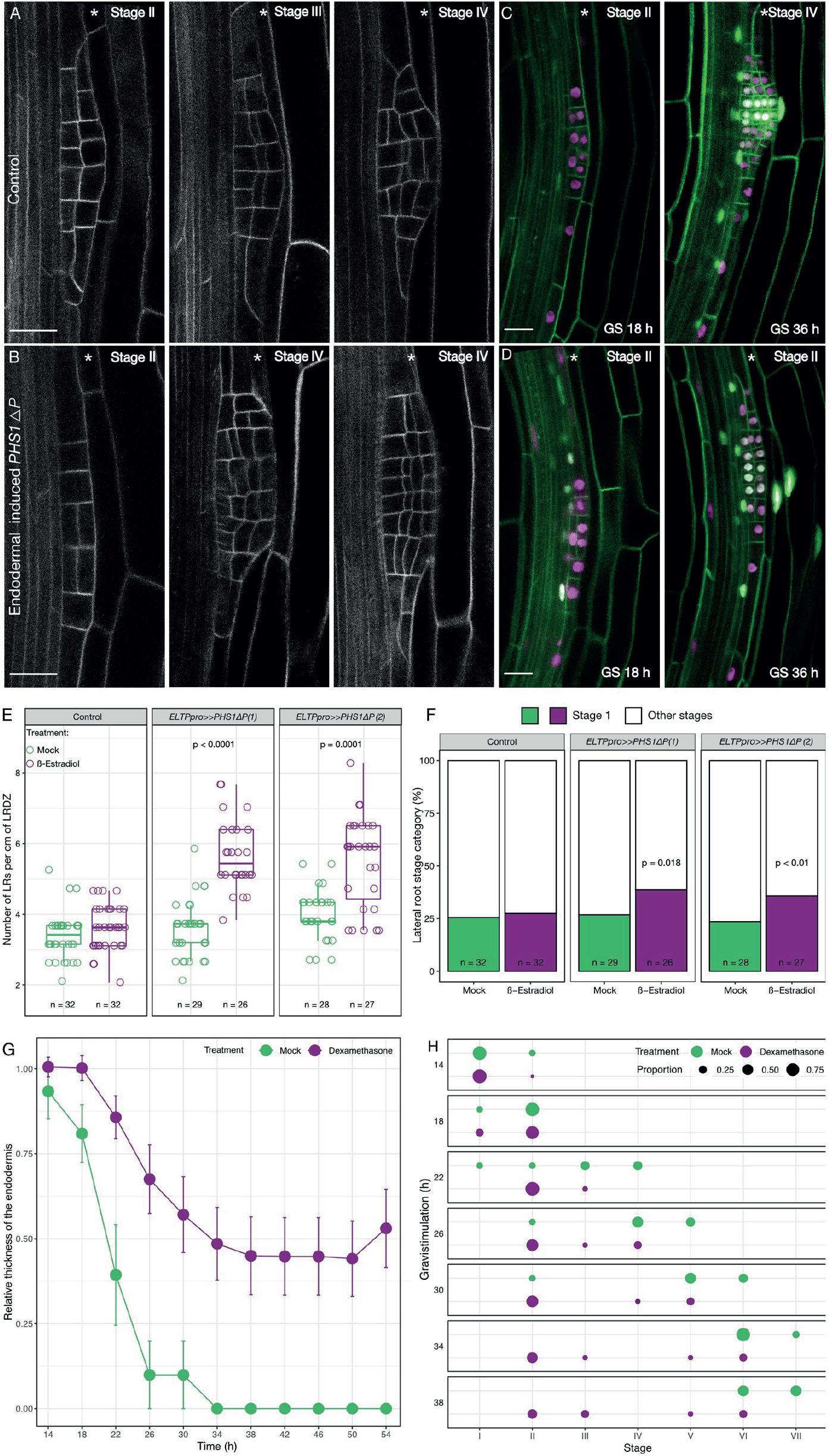
The microtubule cytoskeleton of the endodermis is required for LRP development. (**A-D**) LRP morphology in *ELTPpro>>PHS1*Δ*P* lines visualised by the plasma membrane marker *UBQ10pro::EYFP:NPSN12* (gray) (**A, B**), or *CASP1pro>>PHS1*Δ*P* lines carrying the fluorescent markers *UBQ10pro::GFPx3:PIP14*; *GATA23pro:H2B:3xmCherry*; *DR5v2pro::3xYFP:NLS*; *RPS5Apro::tdTomato:NLS* (sC111) (13) (**C, D**) under mock (**A, C**) or after induction (**B, D**). (**C, D**) Gravistimulation (GS) induced LRP formation observed at 18 h and 36 h after GS. Scale bars = 20 µm. (**E**) LRP density in the LR developmental zone (LRDZ). For each line, Wilcoxon rank-sum test was used to compare LRP densities under the two treatments. (**F**) Analysis of stage I LRP upon disruption of the CMT. Pearson’s Chi-squared test with Yates’ continuity correction was used to assess if LRP distribution is independent of CMT condition. (**G**) Analysis of stage I LRP upon disruption of the CMT. Pearson’s Chi-squared test with Yates’ continuity correction was used to assess if LRP distribution is independent of CMT condition. (**H**) Distribution of LRP developmental stages induced by GS upon induction of *CASP1pro>>PHS1*Δ*P* expression and in mock.

### Endodermal CMT reorganization depends on SHY2-mediated auxin signaling

SHY2-mediated auxin signaling has been shown to control spatial accommodating responses in the endodermis required for LRP formation (14). Accordingly, we investigated whether endodermal, SHY2-mediated auxin signaling was also required for spatially defined CMT domains and their reorganization. We used *CASP1pro::mVenus:MBD* to quantify the response of the endodermal CMT lattice in plants expressing *shy2-2*, a dominant transcriptional repressor of auxin signaling, in the differentiated endodermis (*CASP1pro::shy2-2*) (14). Since LRP formation is blocked in *CASP1pro::shy2-2* mutants, they were treated with indole-3-acetic acid (IAA; 1 µM) to test whether auxin could affect CMT organization in the endodermis. In wild type (WT), 1 µM IAA treatment induces a reorientation of the CMT on the inner side of endodermal cells comparable to that observed upon LR formation. In *CASP1pro::shy2-2* roots, CMTs on the inner side do not reorganize and remain in the same organization as prior to auxin treatment (**Fig. 3A** and **B**). To verify that this effect is specific to SHY2-mediated auxin signaling in the endodermis and not just due to an absence of LRP formation, we repeated the experiment in the slr-1 mutant in which LRP initiation is also blocked, but the SHY2 response in the endodermis is unaffected (24). The reorganization of the CMTs induced by auxin in slr-1 expressing *CASP1pro::mVenus:MBD* is similar to that observed in WT roots (**Fig. 3C**), establishing that SHY2-mediated auxin signaling is directly responsible for the rearrangement of CMTs on the inner side of endodermal cells and is not an indirect consequence of auxin-induced LR development. Interestingly, the reorganization of CMTs on the outer side of WT, *CASP1pro::shy2-2* and *slr-1* is not affected (**Fig. 3C-F**). Consequently, it appears that SHY2-mediated auxin signaling is required for the spatially defined remodeling of CMTs in the endodermis.

**Fig 3.**
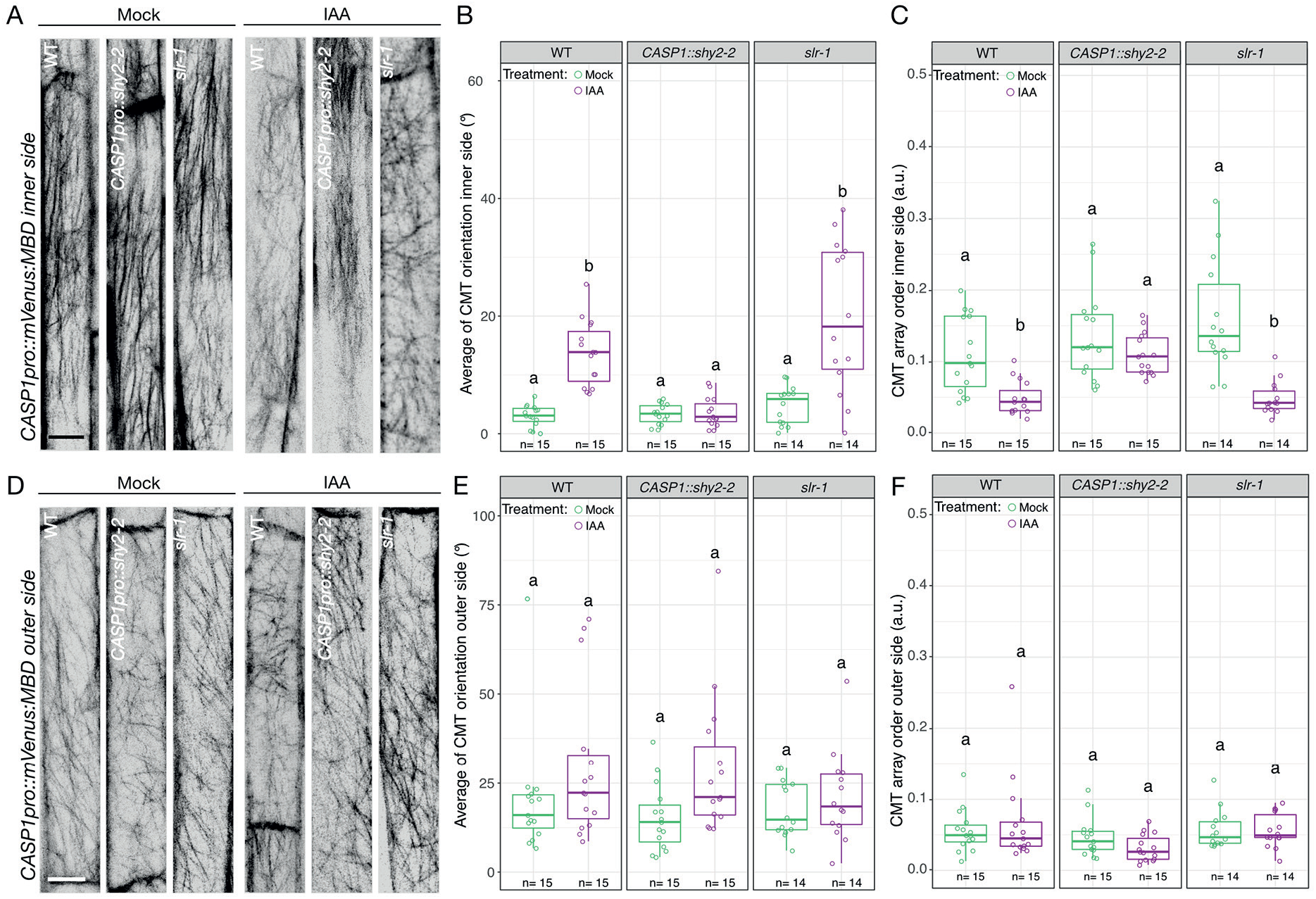
SHY2-mediated local reorganization of CMTs in the endodermis. (**A**) Confocal images of CMT arrays on the inner side of an differentiated endodermis cell in WT, *CASP1pro::shy2-2*, and *slr-1* plants expressing the *CASP1pro::mVenus:MBD* reporter after 24 h of dimethyl sulfoxide (DMSO) or 1 µM Indole-3 acetic acid (IAA) treatment. (**B**) Quantification of CMT orientation (0°–90°, in respect to the long axis of the cell) and (**C**) isotropy (arbitrary units 0-1, 0 = no order, 1 = order) on the inner side of endodermal cells after 24 h of DMSO or IAA (1 µM) treatment. (**D**) Confocal images displaying the organization of CMT arrays on the outer side of an endodermal cell in WT, *CASP1pro::shy2-2*, and *slr-1* plants expressing the *CASP1pro::mVenus:MBD* reporter after 24h of dimethyl sulfoxide (DMSO), or IAA (1 µM) treatment. (**E**) Quantification of CMT orientation (0°–90°, respective to the long axis of the cell) and (**F**) isotropy (arbitrary units 0-1, 0 = no order, 1 = order) on the outer side of endodermal cells after 24 h of DMSO or IAA (1 µM) treatment. Comparison between samples was performed using two-way ANOVA and post hoc multiple comparisons with Tukey’s HSD. Samples with identical letters do not significantly differ (α = 0.05). Scale bar = 10 μm.

### SHY2 regulates MICROTUBULE ASSOCIATED PROTEIN 70-5 in accommodating endodermal cells

We next mined an auxin-induced transcriptome of differentiated endodermal cells for cytoskeleton-related genes that could regulate CMT organization and/ or dynamics (24). We identified *MICROTUBULE ASSOCIATED PROTEIN 70-5* (*MAP70-5*) as a potential SHY2-dependent regulator of CMT organization in the endodermis. Plants expressing a *MAP70-5pro::CITRINE:MAP70-5* fusion (*CITRINE:MAP70-5*) display a spiral or punctate localization pattern in differentiating proto- and metaxylem cells, respectively (**Fig. 4A-C**). This is in agreement with a proposed function of MAP70-5 in regulating secondary cell wall formation during xylem formation [(25, 26) and Supplemental Fig. S3 and Supplemental Movie S1]. We also observed *CITRINE:MAP70-5* expression in the pericycle and endodermis in the early differentiating zone of Arabidopsis roots, the area where lateral root founder cell (LRFC) specification is reported to occur (27) (**Supplemental Fig. S3**). This early expression in the pericycle and endodermis of *MAP70-5:CITRINE* appeared not to be affected in *CASP1pro::shy2-2* roots (**Supplemental Fig. S3**). Interestingly, during LRP formation, we observed induction of CITRINE:MAP70-5 specifically in endodermal cells overlying growing LRP from stage II onwards (**Fig. 4D-L, Supplemental Movie S2**). *MAP70-5* contains a predicted auxin responsive element (TGTCTC) (28, 29) in its promoter region, in line with auxin-dependent regulation. Indeed, auxin treatment induces an expansion of the expression domain of *CITRINE:MAP70-5* in endodermal and cortex cells throughout the root (**Supplemental Fig. S4 A-C**). This auxin-mediated induction of **CITRINE:MAP70-5** is completely blocked in *CASP1pro::shy2-2* seedlings (**Supplemental Fig. S4 D-F**). From this we determine that while endodermal expression of *MAP70-5* can occur independently of auxin activity, the specific induction of *MAP70-5* in endodermal cells overlying the LRP is dependent on SHY2-mediated auxin signaling. Induction of *CITRINE:MAP70-5* in the cortex of *CASP1pro::shy2-2* roots is also blocked, suggesting that this also requires endodermis-mediated auxin signaling. In endodermal cells overlying an LRP, CITRINE:MAP70-5 localizes to filamentous structures in the cell periphery. Four-dimensional live cell imaging of the endodermis during LRP formation revealed that CITRINE:MAP70-5 labels dynamic structures and their organization becomes more isotropic when the LRP reaches stage IV; the point at which it traverses the endodermis (**Fig. 4 N-O** and **Supplemental Movie S3**). We also observed that CITRINE:MAP70-5 displays a differential localization pattern in the accommodating endodermal cells (**Fig. 4 O-Q**). Quantification of signal intensity on the inner and outer side of these cells reveals an enrichment of CITRINE:MAP70-5 (r = 1.44 ± 0.428, n = 25) on the inner side of endodermal cells overlying stages III/IV LRPs (**Fig. 4P**). Imaging of CITRINE:MAP70-5 and a microtubule marker (*ELTPpro::mScarlet-I:MBD*) revealed a partial colocalization between MAP70-5 and CMTs (**Supplemental Fig. S5 A-F**). Depolymerization of CMTs, either by oryzalin or via induction of PHS1ΔP expression in the endodermis (*ELTPpro>>PHS1*Δ*P*), revealed that in both cases, the localization of CITRINE:MAP70-5 gradually shifted from a filamentous to a more diffuse cytosolic localization (**Supplemental Fig. S5 G-L**). Together, these results show that MAP70-5 partially colocalizes with CMTs in endodermal cells and requires intact CMTs for its localization. Further, CITRINE:MAP70-5 preferentially accumulates on the inner side of endodermal cells overlying stage III/IV LRPs, alluding to a potential contribution to the reorganization of CMTs during spatial accommodation.

**Fig 4.**
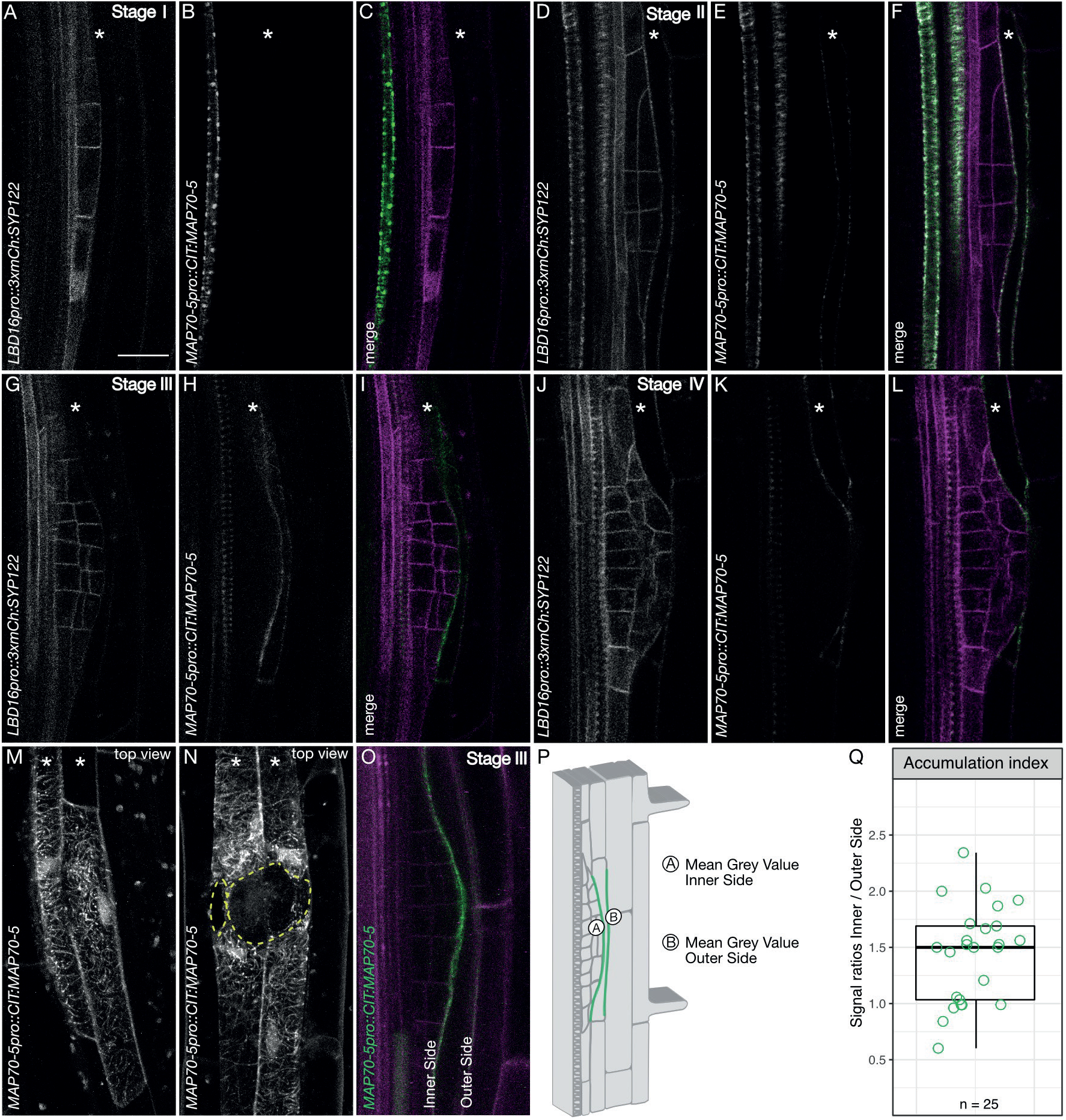
MAP70-5 expression in the endodermis correlates with spatial accommodation. (**A-O**) Confocal images of developing LRs in seedlings expressing *LBD16pro::3xmCherry:SYP122* (plasma membrane, grey (**A, D, G**, and **M**) or magenta (**C, F, I** and **L**) to visualise the LRP and *MAP70-5pro::CITRINE:MAP70-5* (grey; **B, E, H, K** and **M-N** or green; **C, F, I** and **L**). (**A-C**) Stage I: *CITRINE:MAP70-5* is expressed in a differentiating metaxylem cell. (**D-L**) CITRINE:MAP70-5 is induced in endodermal cells overlying the LRP from stage II to IV. (**M-N**) Confocal image stack depicting a surface view of endodermal cells overlying the LRP: (**M**) stage IV, (**N**) stage V. (**O**) Single confocal image showing accumulation of CITRINE:MAP70-5 on the inner side of an endodermal cell overlying a stage IV LRP. (**A-L** and **O**) Images of single confocal sections. (**M-N**) Maximum projections of z-stacks. Stars indicate endodermal cell files. Yellow dashed circles indicate the area of the LRP. (**O**) Stage III LRP showing accumulation of CITRINE:MAP70-5 (green) on the inner side of an overlying endodermal cell. Magenta shows plasma membrane labelled by UBQ10pro::tdTomato:RCI2a (36). (**P**) Schematic representation of inner and outer domains used for quantification of the accumulation index of CITRINE:MAP70-5 in endodermal cells overlying LRP (stages II-IV). (**Q**) Quantification of the mean gray values of the inner and outer domains. For the accumulation index the collected data was normalized and the ratio between the inner a the outer domain was calculated. Scale bar = 20 µm.

### MAP70-5 affects LRP formation in a non-cell autonomous manner

To test whether MAP70-5 is required for reorganization of CMTs during the remodeling of the endodermis throughout LRP formation, we generated *map70-5* mutants, *map70-5-c1* and *map70-5-c2*, which both contain large deletions resulting in an early stop codon (**Supplemental Fig. S6**). Both *map70-5* alleles show a reduction in primary root length and have a smaller LRDZ zone (**Supplemental Fig. S6B-D**). Quantification of the distribution of LRP stages along the LRDZ of both *map70-5* alleles revealed a significant increase in total LRPs (**Fig. 5A**), with both alleles showing a higher proportion of stage I LRPs (**Fig. 5B**), suggesting an increased rate of LR initiation and a delay in emergence. This was confirmed by introgression of the plasma membrane marker *UBQ10pro::EYFP:NPSN12* into map70-5-c1 and comparison of LRP morphology to WT plants (Stages I to IV). In contrast to WT, *map70-5-c1* mutants display flattened LRPs and a turgid endodermis from stage II onward (**Fig. 5 C** and **D, Table S1**). This phenotype is similar to the one observed upon depolymerization of endodermal CMTs (Fig. 2). We then set out to test whether MAP70-5 plays a role in the organization and dynamics of the endodermal CMT lattice during LR formation. Using *CASP1pro::mVenus:MBD* to quantify the organization of the CMT in endodermal cells revealed that the *map70-5-c1* mutant has lost the differential organization of the CMT lattice on the inner side of the cell. In the mutant, the organization of the CMT is comparable to that observed in endodermal cells prior to LRP initiation (**Fig. 5 E-G** and **Fig. 2**), indicating that MAP70-5 is required for the spatial re-organization of CMT arrays in the endodermis. Taken together, these results show that MAP70-5 acts during two phases of spatial accommodation by the endodermis during LRP development. In the first phase, it functions as a negative regulator that confines LRP initiation via CMT array organization on the inner side of endodermal cells. In the second phase, it appears to act as a positive regulator of LRP development and morphogenesis. We propose that MAP70-5 integrates and possibly relays instructive signals via the CMT in the endodermis during LRP formation.

**Fig. 5.**
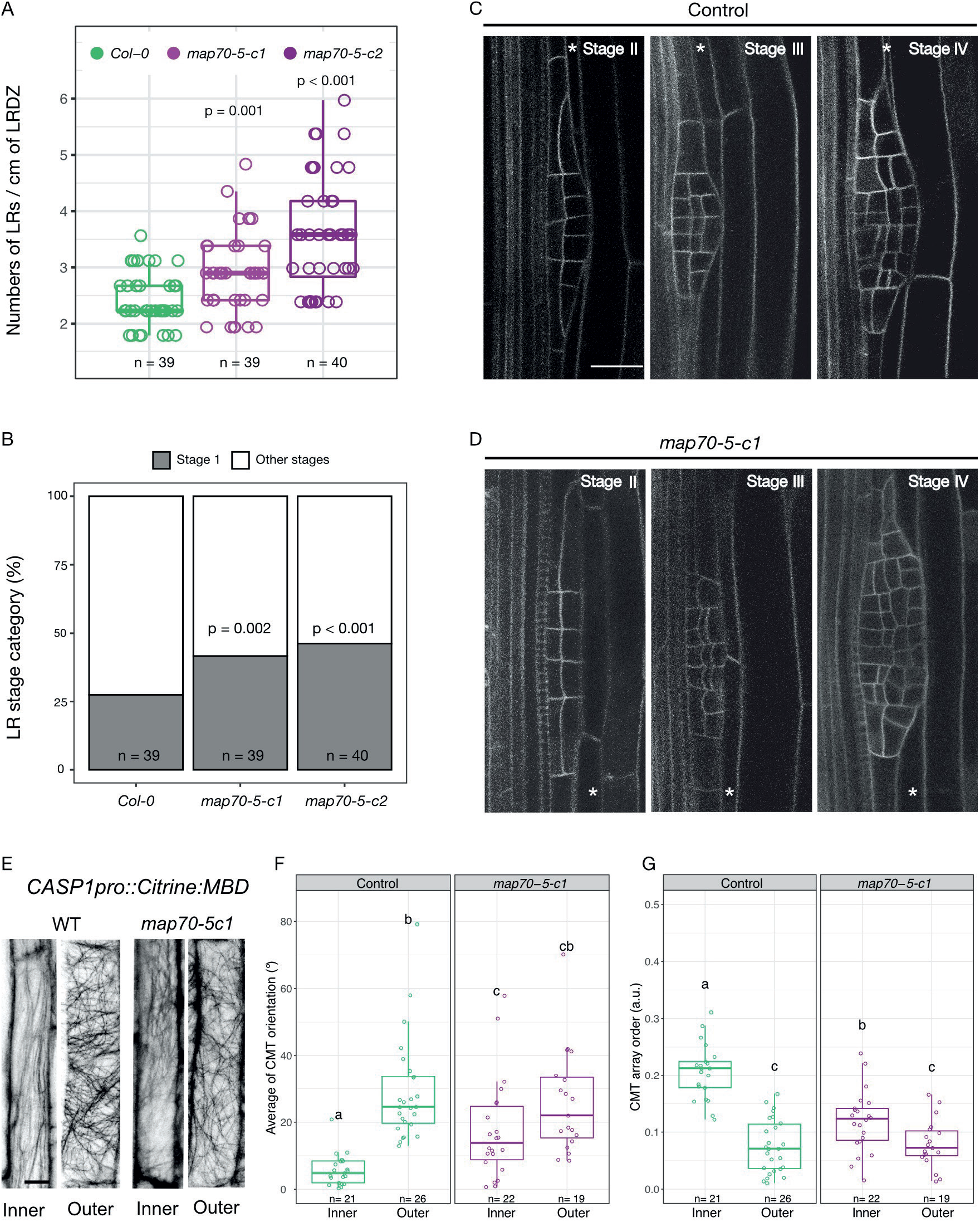
MAP70-5 is required for spatial accommodation responses in the endodermis. (**A**) LR density in the LRDZ of WT compared to *map70-5-c1* and *-c2*. For each line, Wilcoxon rank-sum test with continuity correction was performed to compare LR densities. (**B**) Staging of LRs reveals an accumulation of stage I LRP in the *map70-5-c1* and *-c2* compared to WT roots. Pearson’s Chi-squared test with Yates’ continuity correction was used to assess if LRP distribution is independent in the *map70-5* mutants. (**C** and **D**) LRP morphology in WT and *map70-5-c1* through stages I-IV visualized by the plasma membrane marker UBQ10pro::EYFP:NPSN12. (**E**) Confocal images showing CMTs on the inner and outer side of WT and *map70-5-c1* endodermal cells. (**F**) CMT orientation (0°–90°, respective to the long axis of the cell). (**G**) CMT isotropy on the inner and outer side (arbitrary units 0 = no order, 1 = order). Scale bar = 10 µm. Comparison between samples was performed using two-way ANOVA and post hoc multiple comparisons with Tukey’s test. Samples with identical letters do not significantly differ (α = 0.05).

## DISCUSSION

Plant development, just like in other organisms, relies on the integration of both chemical and physical signals (30-32). In contrast to what we have learned about this process in surface cell layers (33-35), it is still not clear how mechanical conflicts generated during differential growth in deep lying plant tissues are integrated and translated into a developmental output and if similar mechanisms are at play. Previously, we showed that asymmetric CMT organization of LRFCs is required for their asymmetric expansion and to license LRP initiation. Moreover, we showed that endodermal auxin signaling was required for the remodeling of the LRFCs. However, how the endodermis contributed to LRP initiation remained unknown (13).

Here, we identify a molecular mechanism operating in the endodermis that is required to channel organ initiation and development in the XPP. By combining live-cell imaging and cell-type-specific genetic perturbations of microtubule organization, we show that CMTs in the endodermis are required for endodermal remodeling and normal LRP development. CMT arrays on the side of endodermal cells facing the LRFCs are more anisotropic than those on the other side of the same cell, and this is required for proper LRP formation. To accommodate the newly formed LRP, the CMTs on the inner side of overlying endodermal cells will subsequently become more isotropic to facilitate thinning of endodermal cells. However, the observation that depolymerization of endodermal CMTs, which is assumed to make cells less mechanically resistant, results in an increase in stage I LRPs and interferes with endodermal thinning (**Fig. 2**). This rather suggests that a tight regulation of endodermal CMT organization is required to coordinate the expansion growth of the LRP. Initially, anisotropic CMT arrays on the inner side of overlying endodermal cells are required to suppress LRP initiation, possibly by providing mechanical feedback. After initiation, the endodermal CMT arrays need to become more isotropic in order to channel the outgrowth of the LRP. Thus, a regulated switch from anisotropic to more isotropic endodermal CMT arrays is necessary for both the perception of a signal instructing the endodermis to remodel to initiate as well as for the completion of this developmental program and both processes require a SHY2-dependent auxin signaling module.

Analyzing the SHY2-regulated, auxin-mediated transcriptome of endodermal cells (24), we identified MAP70-5 as a candidate regulator of CMT organization during spatial accommodation. *CITRINE:MAP70-5* is induced in overlying endodermal cells, and partially colocalizes with CMTs, whereas *map70-5* mutants show an increase in stage I LRP and altered morphogenesis up to stage IV LRP (**Fig. 5**). Endodermis-specific CMT depolymerization and the *map70-5* mutants have similar phenotypes, suggesting that MAP70-5 is an important relay in the communication between the LRP and the endodermis. In *map70-5* mutants, the endodermis is predicted to be altered in its ability to perceive or respond to the outgrowth of the LRP. Quantification of the CMT array order on the inner and outer side of endodermal cells revealed that MAP70-5 also functions before LRP formation. Endodermal cells in roots of *map70-5* mutants no longer display the spatially defined, differential CMT array order (**Fig. 5 E-G**). It appears that CMT arrays on the inner side are already in a configuration that facilitates LR initiation. A plausible mechanism is that MAP70-5 locally regulates CMT array order on the inner side of the endodermis via bundling, and this is a part of a mechanism that spatially restricts LRP initiation. Subsequently, MAP70-5 is required for proper endodermal thinning to properly channel LRP organogenesis, since *map70-5 mutants* also display a significant increase in flattened LRP, which we also observed upon depolymerization of endodermal CMTs. This is again in support that CMT organization needs to be tightly regulated, in order accommodate organ growth.

How might MAP70-5 regulate the cell shape of endodermal cells overlying LR founder cells? Since these endodermal cells are fully differentiated, it is unlikely that the observed effects are due to defects in the guidance or dynamics of the cell wall synthesis machinery. The fact that endodermal cells of map70-5-c1 roots have lost the differentially organized CMT array order on the inner side would support a direct role for the microtubule cytoskeleton itself during founder cell - endodermis communication and possibly also during LRP outgrowth. Moreover, the punctate localization of CITRINE:MAP70-5 at the cell periphery suggests that MAP70-5 might be partially localized at the plasma membrane. Since the overlying endodermal cells must undergo a drastic change in cell volume to accommodate the outgrowth of the LRP, we hypothesize that MAP70-5 is required to transduce the mechanical constraints detected at the interface between the LRP and neighboring endodermal cells and to regulate CMT array order to channel organogenesis. Thus, we propose a role for MAP70-5 as an integrator of mechanical constraints generated during the re-establishment of 3D-differential growth within a tissue.

### Experimental Procedures

## MATERIALS AND METHODS

### Plant materials and manipulation

The *Arabidopsis thaliana* Columbia ecotype (Col-0) was used. Seeds were surface sterilized (Sodium hypochlorite 5 % and 0.01 % Tween 20 or Ethanol 70 % and SDS 0.1 %) and placed on ½ Murashige and Skoog (MS) medium containing 1 % agar (Applichem or Duchefa). Following stratification (4°C in the dark, >t 24 h), seedlings were grown at 22°C vertically under constant light or long day conditions (16 h light / 8 h dark). Agrobacterium tumefaciens (GV3110) based plant transformation was carried out using the floral dip method (1). All plant lines examined were homozygous if not indicated otherwise. Homozygosity was either determined upon the presence of the Fast Red cassette, by antibiotic resistance, verification of the fluorescent fusion proteins at the microscope and/ or PCR. Besides the below mentioned created vectors and plants lines, this study uses the triple marker line *UBQ10pro::GFP:PIP1* x *GATA23pro::H2B:Mcherry* x *DR5v2pro::NLS-3xYFP* sC111 (2) and the membrane marker line *UBQ10pro::EYFP:NPSN12* (3). For experiments with inducible gene expression and/or drug treatments, β-estradiol (β-est), dexamethasone (Dex) and oryzalin stocks, were dissolved in EtOH, DMSO or water, were used as indicated.

### Construction of vectors and transformation

To generate *ELTPpro>>PHS1*Δ*P* (*pFR-ELTPpro-XVE>>PHS1*Δ*P*), *pEN-4_ELTPpro-XVE_1R, pEN-1_ PHS1*Δ*P_2* and *pEN-2r_mCherry_3* were recombined into *pFastRed-3xGW* (4). To generate *CASP1pro>>PHS1*Δ*P* (*CASP1pro::LhG4-GR-6xOP:PHS1*Δ*P-mCherry*), we used GreenGate assembly (5) to combine *pGGM-CASP1pro::LhG4-GR* (6) with *pGGN-6xOP:PHS1*Δ*P-mCherry-FastRed* (2).

To generate *pGr179-CASP1pro::mVENUS:MBD*, KpnI was used to exchange *XPPpro* with *CASP1pro* in *pGr179-XPPpro::mVENUS:MBD* (2) from *CASP1pro::CITRINE:SYP122* (7) using KpnI. To generate *pGr179-LBD16pro::3xmCherry:SYP122*, KpnI was used to exchange *XPPpro* with *LBD16pro* in *pGr179-XPPpro::3xmCherry:SYP122* (4). To generate *pUC57-L4_ MAP70-5pro_R1*, 1573bp of upstream sequence of *MAP70-5* (AT4G17220) was amplified and cloned into KpnI-digested *pUC57-L4_KpnI/XmaI_R1*. For *pEN-2R_MAP70-5_3*, the genomic sequence of *MAP70-5* (as amplified and recombined into pENTR1-2 using BP CLONASE II (www.thermofisher.com). To assemble *MAP70-5pro:CITRINE:MAP70-5, pUC57-L4_MAP70-5pro_R1, pEN-1_CITRINE_2* and *pEN-2R_MAP70-5_3* were recombined into *pH7m34GW,0*. To generate *pFR-ELTPpro::mScarlet-I:MBD, pEN-4_ELTPpro_R1, pEN-1_mScarlet-I_2* and *pEN-2R_MBD_3* was recombined into *pFastRed-3xGW. pEN-1_mScarlet-I_2* was amplified and recombined into pENTR1-2 and *pEN-2R_MBD_3* was synthesized (www.thermofisher.com). Expression vectors were assembled using MultiSite Gateway Cloning with LR CLONASE II Plus (www.thermofisher.com). For CRISPR/Cas9-mediated generation of MAP70-5 mutants, we used *pFR-UBQ-CAS9-1xGW* (8). Three different sgRNA constructs were cloned, targeting different areas of MAP70-5, using a combination of GreenGate (5) and Gateway cloning. Primers for amplifying fragments used for cloning and sgRNAs used for generating *map70-5-c1* and *-c2* mutants are shown in **Table S2** and **Supplemental Fig. S7**. In addition to the deletion shown in **Supplemental Fig. S6**, *map70-5-c1* also has an insertion of a single A after position 1774 at the target site of sgRNA3-1. This is downstream of the early stop codon introduced by the large deletion. For transient expression in *Nicotiana benthamiana, pFR-35Spro::mScarlet-I:MBD, pFR-35Spro::mScarlet-I:MAP70-5, pFR-35Spro::mCITRINE:map70-5c1* and *pFR-35Spro::mCITRINE:map70-5-c2* were generated and plants were infiltrated as described (9). The genomic sequence of *map70-5-c1* and *map70-5-c2* were amplified and recombined into pDONR1-2. Resulting pENTRY vectors were assembled into *pFastRed-3xGW*: *pEN-4_35spro_R1, pEN-1_mScarlet-I_2* and *pEN-2R_MBD_3*; *pEN-4_35spro_R1, pEN-1_mScarlet-I_2* and *pEN-2R_ MAP70-5*; *pEN-4_35spro_R1, pEN-1_mCITRINE_2* and *pEN-2R_map70-5-c1_3*, and *pEN-4_35spro_R1, pEN-1_mCITRINE_2* and *pEN-2R_map70-5-c2_3*. 35S promoter driven expression of *mCitrine:map70-5-c1* and of *mCitrine:map70-5-c2* alleles resulted only faint cytosolic fluorescence in the cytosol of infiltrated Tobacco leaves (**Supplemental Fig. S6**).

### Microscopy

Live cell imaging performed with a Leica TCS SP8X-MP, Leica TCS SP8X, or a Leica TCS SP8-Dive system, equipped with a resonant scanner (8 kHz), a 63x, NA = 1.2 water immersion objective, a 63x, NA = 1.3 glycerol immersion objective, a 63x, NA = 1.4 oil immersion objective or a 40x, NA = 1.2 water immersion objective. For excitation of CITRINE, mVENUS and mCherry either an insight DS+ Dual ultrafast NIR laser or a Chameleon Vision II tuned to 960 nm was used for multiphoton excitation. For detection, non-descanned super-sensitive photon-counting hybrid detectors (HyD), operated in photon-counting mode, were used. EYFP/ mVENUS fluorescence was filtered with a CFP/YFP filter cube (483/32 & 535/30 with SP) or using the 4-Tune detector set to the same detection window. For colocalization of MAP70-5pro::CITRINE:MAP70-5 and ELTPpro::mScarlet-I:MBD, we used a white light laser (WLL) for excitation (517 nm for Citrine and 561 nm for mScarlet-I). Fluorescence was detected using HyD detectors operating in photon-counting mode (527-560 nm for CITRINE and 575-620 nm for mScarlet-I. For CMT array observations, CASP1pro::mVenus:MBD was excited with a WLL at 514nm and detected using the HyD detector in standard mode (525-560 nm). Images were acquired using sequential scanning mode to minimise cross-talk between different detector channels. The acquired z-stacks in this study were taken in step sizes ranging from 0.5-1.0 µm. For long-term imaging, time-lapse data were acquired every 15 - 30 min for up to 16 h (Supplemental Movies S1-3).

### LRP analysis

For endodermal specific MT disruption, *ELTPpro>>PHS1*Δ*P* in the *UBQ10pro::EYFP:NPSN12* (W131Y;(3)) background, and *UBQ10pro::EYFP:NPSN12* as control lines were germinated on plates containing either 5 µM ß-est, or EtOH (mock). For the *CASPpro>>PHS1*Δ*P* in the *UBQ10pro::GFPx3:PIP14*; *GATA23pro:H2B:3xmCherry*; *DR5v2pro::3xYFP:NLS*; *RPS5Apro::tdTomato:NLS* (sC111) background (2), plants were germinated on plates for 5 days then shifted to plates containing either 10 µM dexamethasone, or ethanol (mock) and gravistimulated by rotating the plate 180°.

Phenotyping and staging along the LRDZ was performed by live cell imaging (Leica TCS SP8X-MP). Experiments were repeated six times. To quantify LRP in the map70-5 mutants along the LRDZ, seedlings were cleared as published (10), and staged using a Nikon E800 upright microscope equipped with differential interference contrast. For visualization of the cell outline, *map70-5-c1* plants were crossed with the W131Y cell membrane marker line and homozygous F3 plants imaged and staged from stage I until stage IV (Leica SP8-MP DIVE 960 nm excitation). To determine total root and LRDZ length of *ELTPpro>>PHS1*Δ*P* in the *UBQ10pro::EYFP:NPSN12* and *map70-5* mutants, seven day old seedlings were scanned and either measured from the root tip to the first visible emerged LR (LRDZ) or from the root tip to the hypocotyl (total root length), using-the segmented line tool in FIJI. To analyze differentially expressed *CITRINE:MAP70-5* in accommodating endodermal cells overlying the LRP, a combination of natural- and gravistimulus-induced LRs were imaged (Leica TCS SP8X-MP). For the former, plants were grown for 6 days. For the latter, seedlings were turned 90° after 5 days, and imaged after a further 24 h. Using the segmented line tool in FIJI, ratios between mean intensity values of CITRINE:MAP70-5 inner and outer side were calculated, describing the accumulation index (Fig. 2 O and P).

### Image processing and assembly

The acquired microscopy images were processed with FIJI (v2.0.0-rc-59/1.51k, https://fiji.sc/) and Affinity Photo v1.6.7. Figure assembly was performed with Affinity Designer v1.83. and Adobe Illustrator 2021 (V25.0).

### Data visualization and statistics

All experiments were performed at least three times. Sample size (n) for each plant line and treatment are denoted in the figures. Statistical comparisons (Wilcoxon rank-sum test, Pearson’s Chi-squared test), box- and bar plots were made with R software (11). Methods and p-values are summarized in figure legends.

### IAA treatments

To examine endodermal microtubule organization upon IAA treatment in different genetic backgrounds (**Fig. 3**), five-day old seedlings were transferred to IAA containing medium (1 µM) for 21 h, and subsequently imaged (Leica TCS SP8X). To ensure the same developmental stage was studied, only endodermal cells adjacent to the fifth cortex cell above the first differentiated protoxylem cell were imaged from the top. From this, z-stacks were generated. During imaging, the seedlings were kept in microscopy chambers and covered with a thin layer of medium either supplemented with IAA or mock. By the end of the experiment, seedlings had been exposed to IAA for 24 h. To test whether MAP70-5 expression is dependent on auxin signaling (Supplemental Figure 6), five-day old seedlings were transferred to IAA containing liquid medium (10 µM) and imaged after 24 h (Leica TCS SP8 MP).

### CMT array analysis

The Fiji plugin FibrilTool (12) was used to analyze CMT orientation and CMT order in the endodermis. Z-stacks of endodermal cells expressing CASP1pro:mVenus:MBD were acquired (Leica TCS SP8X), either on the inner or outer side of the endodermal cell. Maximum projections of the acquired z-stacks were generated and aligned (via rotation) to the longitudinal axis of cells, resembling 0°. Regions of interest (ROIs) were set in order to analyze the CMT orientation, and absolute values (from 0° to 90°) were used to visualize the direction of CMTs. The anisotropy analysis describes the microtubule organization in arbitrary units (a.u.). 0 indicates isotropic- (not-ordered), and 1 anisotropic (ordered) CMT organization.

## ACKNOWLEDGEMENTS

We thank Niko Geldner for providing insightful feedback on the manuscript and the Center of Microcopy and Image Analysis of the University of Zurich for excellent service and support. Funding: Work in the Vermeer laboratory was supported by grants from the Swiss National Science Foundation (Schweizerischer Nationalfonds zur Förderung der Wissenschaftlichen Forschung; PP00P3_157524, 316030_164086 and 310030_197568), the Netherlands Organization for Scientific Research (Nederlandse Organisatie voor Wetenschappelijk Onderzoek; NWO 864.13.008) and support from the University of Neuchâtel. D.S. received support from a Forschungskredit from the University of Zürich (FK19-112). Work in the Maizel lab was supported by Work was supported by the Deutsche Forschungsgemeinschaft FOR2581, the Boehringer Ingelheim Foundation. B.J.R.H. was supported by the Consejo Nacional de Ciencia y Tecnología of Mexico (CONACyT, grants 769058 and 740701).

## Author contributions

A.M. and J.E.M.V. conceived, designed and coordinated the project. D.S., B.J.R.H., A.V.B., M.N., Z.W., S.M.M., J.E.M.V. designed and performed experimental work. R.U. and S.F. provided new reagents. D.S., B.J.R.H., A.M. and J.E.M.V. wrote the manuscript and all authors were involved in the discussion of the work.

## Competing interests

Authors declare no competing interests.

## Data and materials availability

No restrictions are placed on materials, such as materials transfer agreements. Details of all data, code, and materials used in the analysis are available in the main text or in the supplementary materials.

## SUPPLEMENTAL MATERIALS

**Fig. S1.**
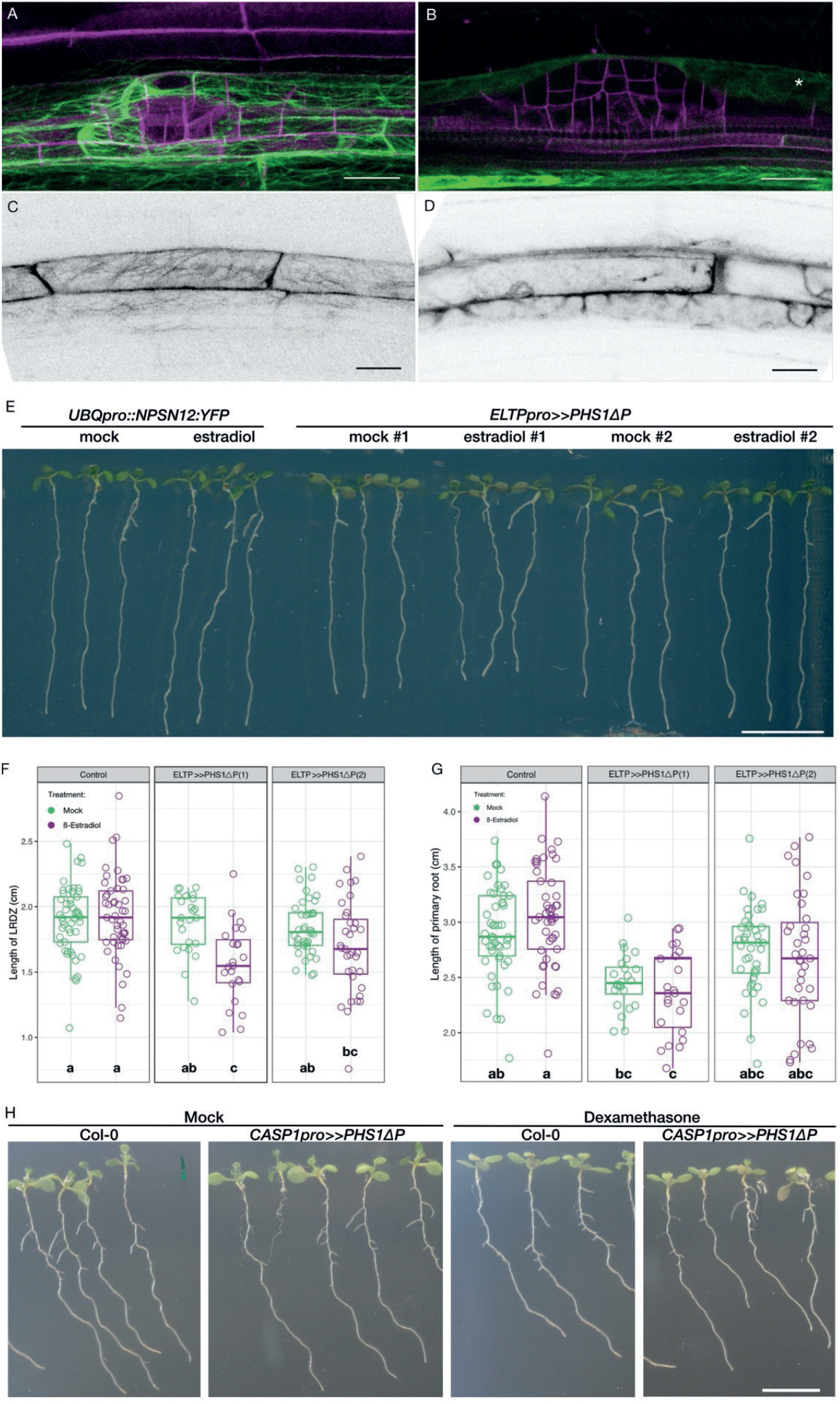
Interfering with endodermal CMT organization impacts LR formation. (**A**) CASP1pro::mVenus:MBD (green) labelling microtubules, and LBD16pro::3xmCherry:SYP122 (magenta) the plasma membrane in the LRP, under mock conditions. (**B**) Disruption of the CMT in the endodermis after *ELTPpro>>PHS1*Δ*P* induction (5 µM ß-est, 24 h). (**C**) *CASP1pro>>PHS1*Δ*P / CASP1pro::mVenus:MBD* showing CASP1pro::mVenus:MBD under mock (H_2_O) or (**D**) upon dexamethasone (Dex) treatment (10 µM). Scale bar 20 µm. (**E**) Image of seven day old seedlings carrying *UBQ10pro::EYFP:NPSN12* and *ELTPpro-XVE>>PHS1*Δ*P* grown on either 5 µM β-est (estradiol) or mock (EtOH) supplemented medium. Scale bar = 1 cm. (**F**) Quantification of the LRDZ and (**G**) total root length. Statistical analysis was performed using one-way ANOVA followed by post-hoc multiple comparisons with Tukey’s test. Samples with identical letters do not significantly differ (α = 0.05). (**H**) Eight-day old seedlings of marker lines sC111 (Col-0) and sC111 combined with *CASP1pro>>PHS1*Δ*P* on mock (H_2_O) or Dex-supplemented medium (10 µM).

**Fig. S2.**
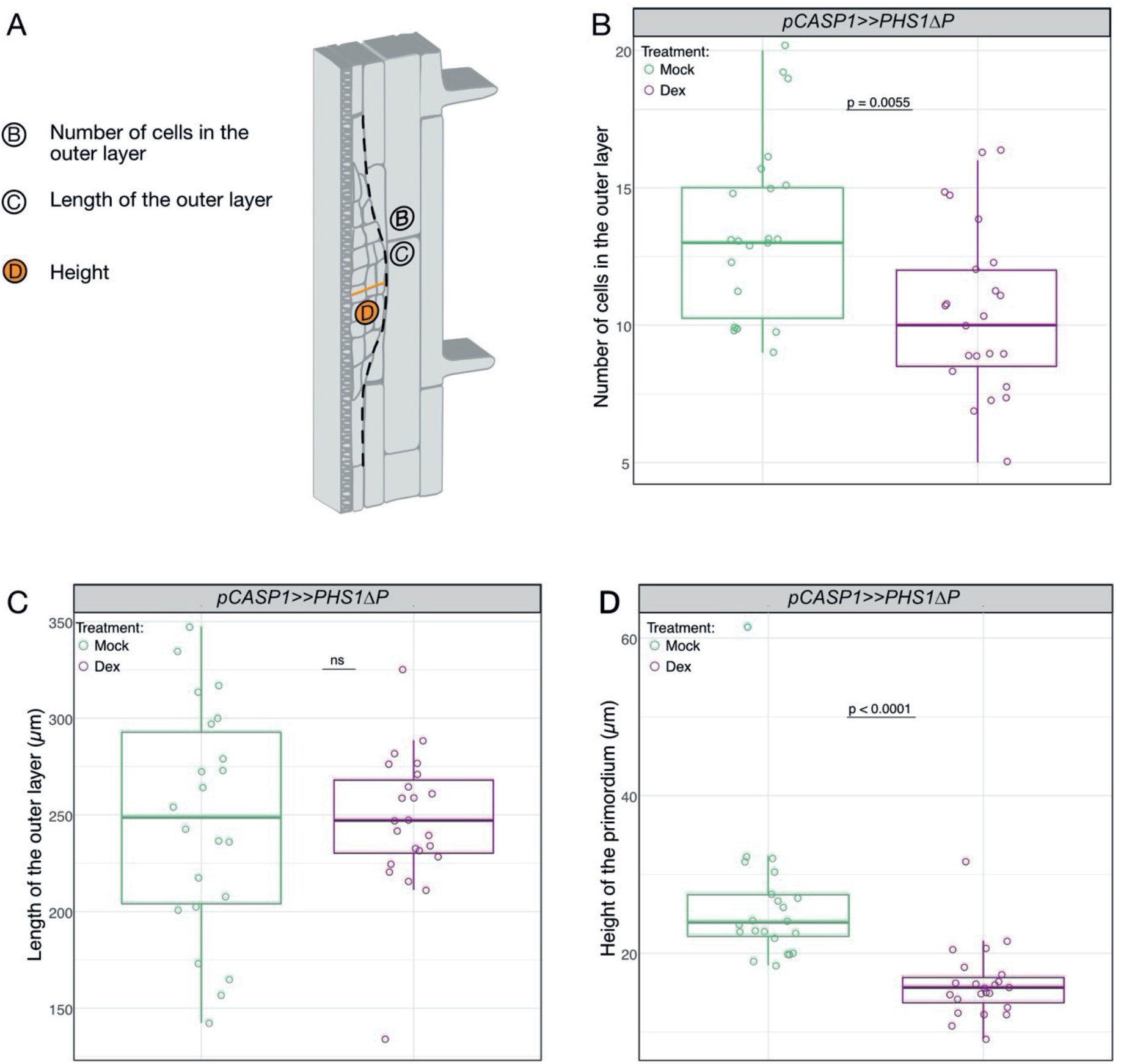
Quantification of the LR morphology after 36 h gravistimulation. (**A**) Drawing of a longitudinal root section displaying a stage III LRP. Letters in (**A**) indicate data represented in (**B-D**). (**B**) Number of cells in the outer layer of the LRP, (**C**) length of the outer layer of the LRP in µm, and (**D**) maximum height of the LRP. The dashed line flanks the outline of the LRP and displays where the measurements were taken and cells were counted. Statistical analysis of *CASP1pro>>PHS1*Δ*P* in the sC111 background on mock (H_2_O, n = 22) and (Dex, n = 23) was performed via a Wilcoxon rank-sum test with continuity correction.

**Fig. S3.**
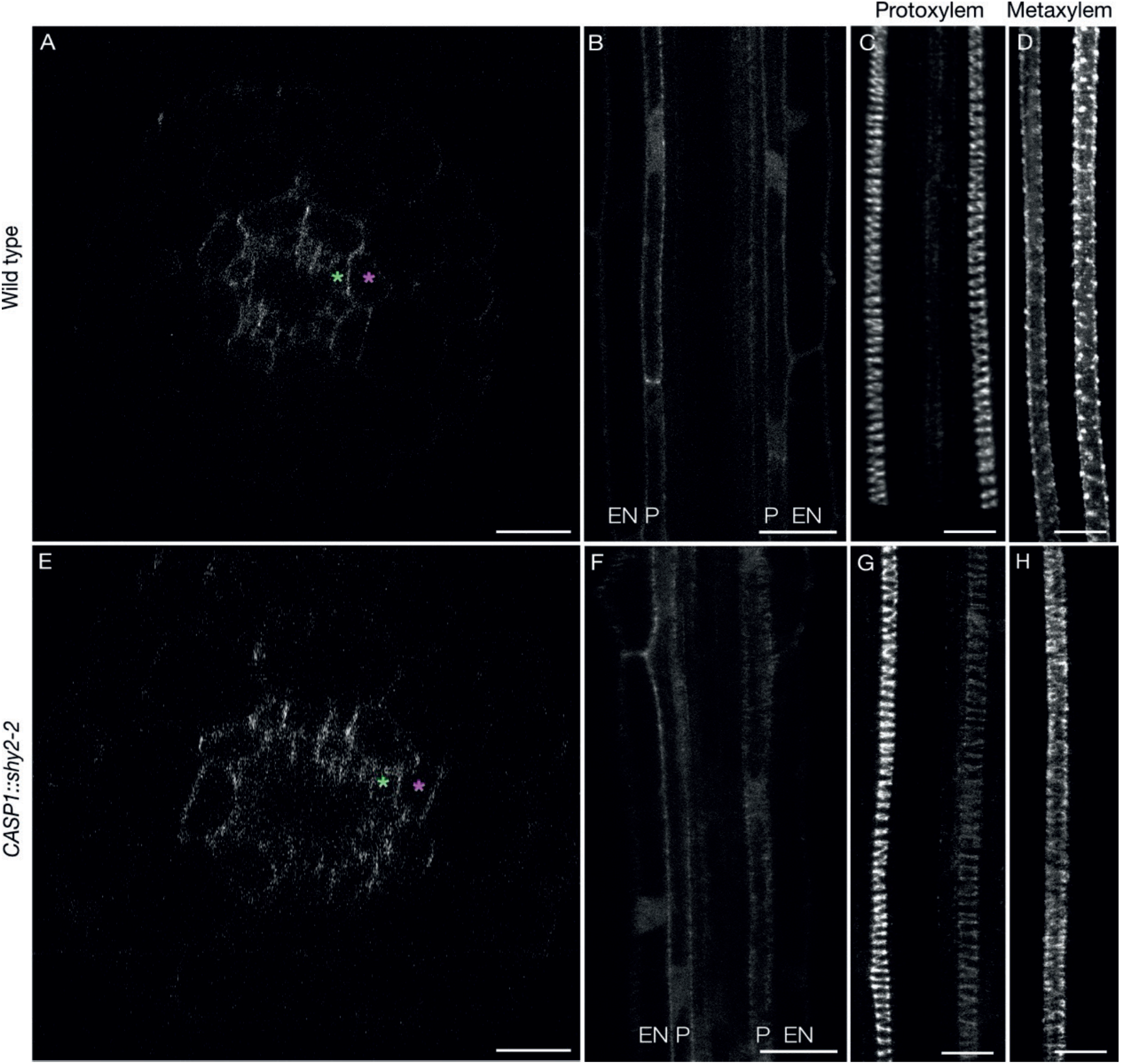
*MAP70-5pro::CITRINE:MAP70-5* is expressed in different root cell types. Localization of MAP70-5pro::CITRINE:MAP70-5 in the root of a seven day old seedling. (**A, B**) In the root hair development zone, *MAP70-5pro::CITRINE:MAP70-5* is expressed in the pericycle and endodermis. (**C, D**) *MAP70-5pro::CITRINE:MAP70-5* is also expressed in isolated, differentiating proto- and meta-xylem cells, respectively. (**E, F**) *MAP70-5pro::CITRINE:MAP70-5* is expressed in the pericycle and endodermis in the root hair development zone in the *CASP1pro::shy2-2* seedlings. (**G, H**) *MAP70-5pro::CITRINE:MAP70-5* expression in isolated differentiating proto- and metaxylem cells in *CASP1pro::shy2-2* seedlings. (**A, E**) Orthogonal view of a root in the early differentiation zone. Green asterisks indicate the pericycle and magenta asterisks mark the endodermis. (**A, E**) Cross sections extracted from confocal z-stacks. (**C, D, G-H**) Maximum projection of a z-stack. Scale bars 10 µm (**C, D, G, H**) and 20 µm (**A, B, E, F**). P = pericycle, EN = endodermis.

**Fig. S4.**
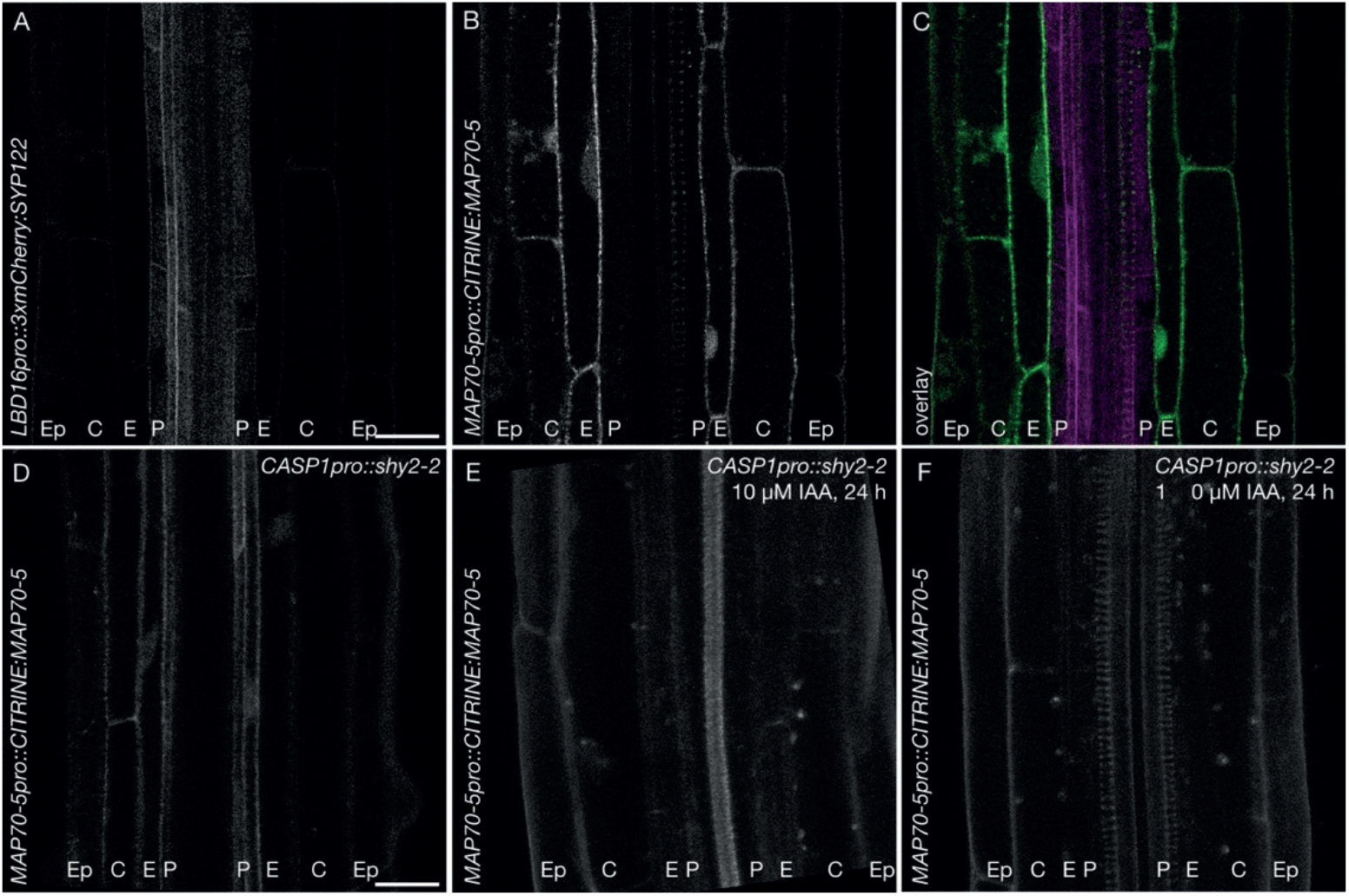
Endodermal *MAP70-5* expression depends on SHY2 mediated auxin signaling. (**A-C**) Confocal images of seedlings co-expressing the plasma membrane marker *LBD16pro::3xmCherry:SYP122* (grey (**A**), magenta (**C**)). (**B**), and MAP70-5pro::CITRINE:MAP70-5 (grey (**B**), green (**C**)) after auxin treatment (10 µM IAA, 24 h) and overlay of both channels (**C**). (**D**) *MAP70-5pro::CITRINE:MAP70-5* expression in *CASP1pro::shy2-2* in the early elongation zone where the *CASP1pro* is not active yet. (**E**) Auxin-mediated induction of MAP70-5pro::CITRINE:MAP70-5 in the endodermis and the cortex is blocked in *CASP1pro::shy2-2* plants. (**F**) The expression of *CITRINE:MAP70-5* in the protoxylem is not affected in *CASP1pro::shy2-2* and IAA treatment (10µM, 24hr). Scale bar is 20 µm. P = pericycle, E = endodermis, C = cortex, Ep = epidermis.

**Fig. S5.**
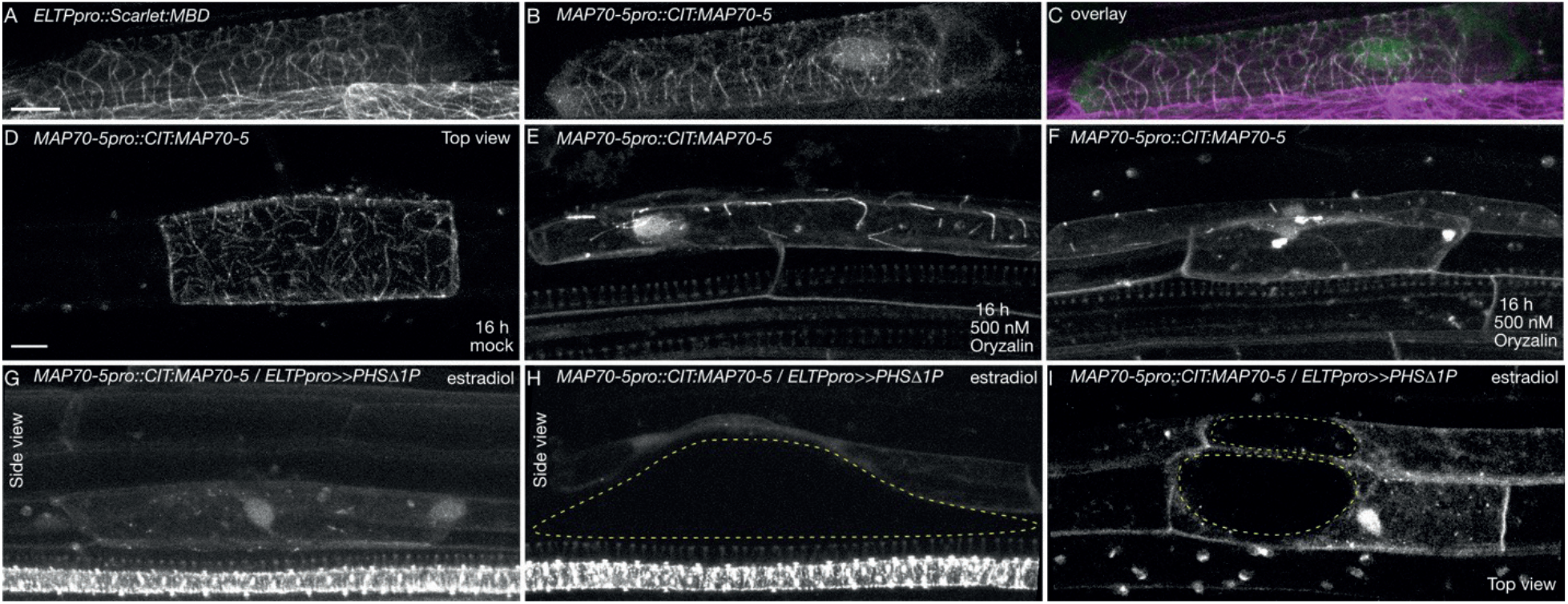
MAP70-5 partially colocalizes with microtubules and depends on an intact CMT cytoskeleton for correct subcellular localization. (**A, C**) Confocal images of endodermal cells co-expressing *ELTPpro::mScarlet-I:MBD* (grey (**A**), magenta (**C**)) and (**B, C**) MAP70-5pro::CITRINE:MAP70-5 (grey (**B**), green (**C**)) showing partial colocalization. (**D-F**) CITRINE:MAP70-5 localization after 16h mock (**G**) or oryzalin treatment (500 nM, **H** and **I**). (**G**) Stage III/IV, (**H**) stage II/III, and (**I**) stage III/IV LRP. (**J-K**) Disruption of the CMT in the endodermis after *ELTPpro>>PHS1*Δ*P* induction in *MAP70-5pro::CITRINE:MAP70-5* expressing lines. (**J**) Stage II/III, (**K**) Stage III/IV, and (**L**) Stage IV/V. All images are maximum projections of confocal z-stacks. Scale bar 10 µm.

**Fig. S6.**
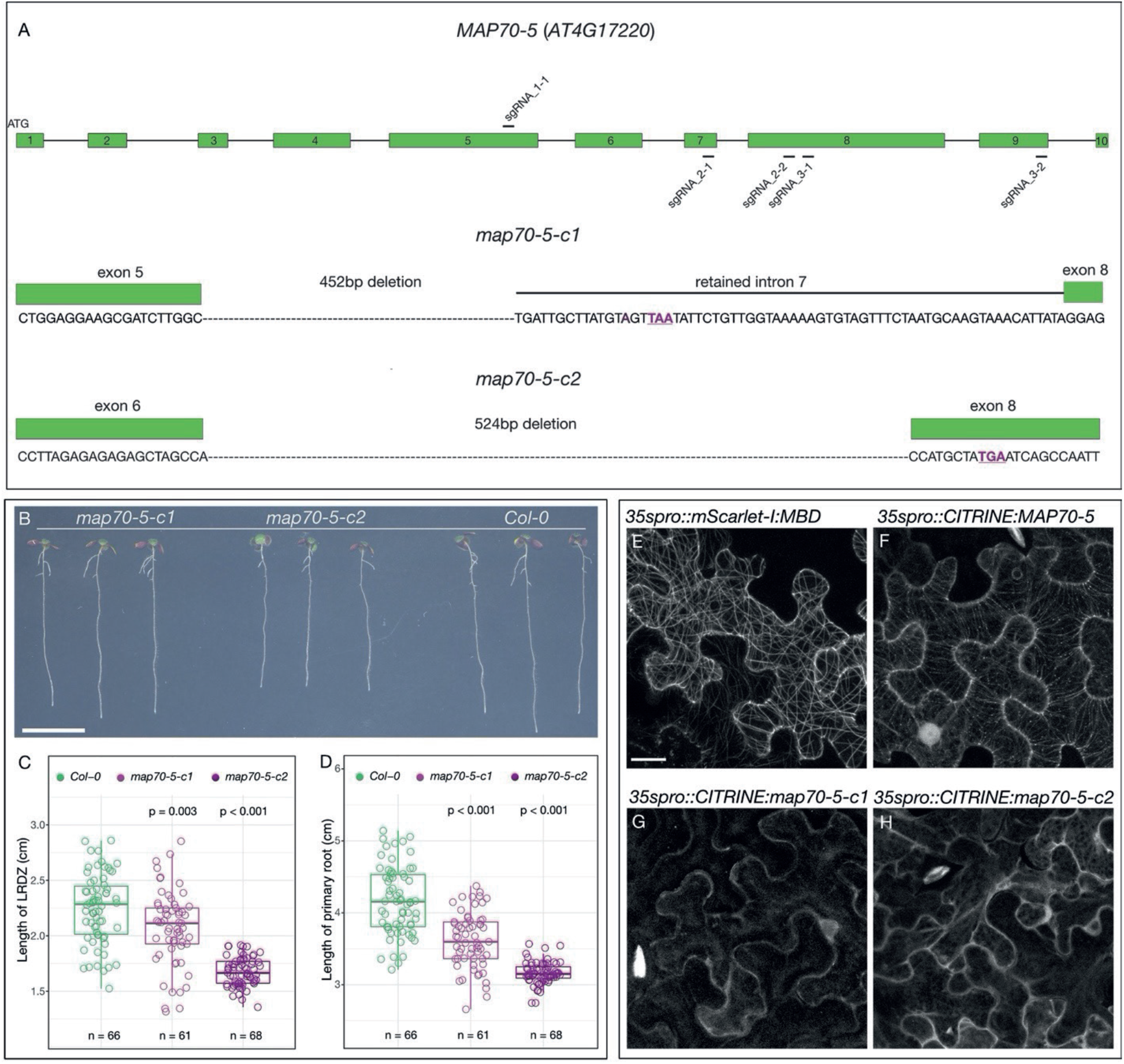
*map70-5* mutants. (**A**) Schematic representation of the gene model of *MAP70-5* and the position of the different sgRNAs used to generate *map70-5* mutants. Bottom panel shows generated deletions that all result in early stop codons, as highlighted in underlined purple letters. (**B**) Image of seven-day old *map70-50-c2, map70-50-c1* and *Col-0* seedlings. Scale bar 1 cm (**C**) Quantification of the LRDZ length and (**D**) of the total root length. Both CRISPR alleles are significantly shorter than the wild type. (**C, D**) Two separate student’s t-tests on Col-0 and *map70-50-c1* and Col-0 and *map70-50-c2* were performed. Transient ectopic expression of *35Spro::mScarlet-I:MBD* (**E**), *35Spro:CITRINE:MAP70-5* (**F**) shows labelling of CMTs, whereas transient ectopic expression of *35Spro::CITRINE:map70-5-c1* (**G**) or *35Spro::CITRINE:map70-5-c2* (**H**) results in faint cytosolic fluorescence.

---

**Supplemental Table S1.** Quantification of LR morphology changes in the *map70-5-c1* mutant.

| Primordia | Stage I |  | Stage II |  | Stage III |  | Stage IV |  |
| --- | --- | --- | --- | --- | --- | --- | --- | --- |
| Observations | Wild type | Aberrant | Wild type | Aberrant | Wild type | Aberrant | Wild type | Aberrant |
| Control (15 roots) | 28 (100%) | 0 (0%) | 26 (76.47%) | 8 (23.53%) | 8 (75%) | 2 (25%) | 10 (100%) | 0 (0%) |
| *map70-5c1* (28 roots) | 56 (85.71%) | 9 (14.9%) | 17 (23.97%) | 57 (77.03%) | 7 (33.3%) | 174 (66.6%) | 6 (76.09%) | 17 (73.91%) |

Observations of mis-shaped primordia and cell division pattern-altered LRPs, and/or delay in endodermis thinning (absolute numbers and percentage). Six to seven day old seedlings were staged until the first primordium traversed the endodermis (mainly stage IV in Control) For visualization of the cell outline, both Control and *map70-5-c1* carry the plasma membrane marker *UBQ10pro::EYFP:NPSN12*.

**Supplemental Table S2.**
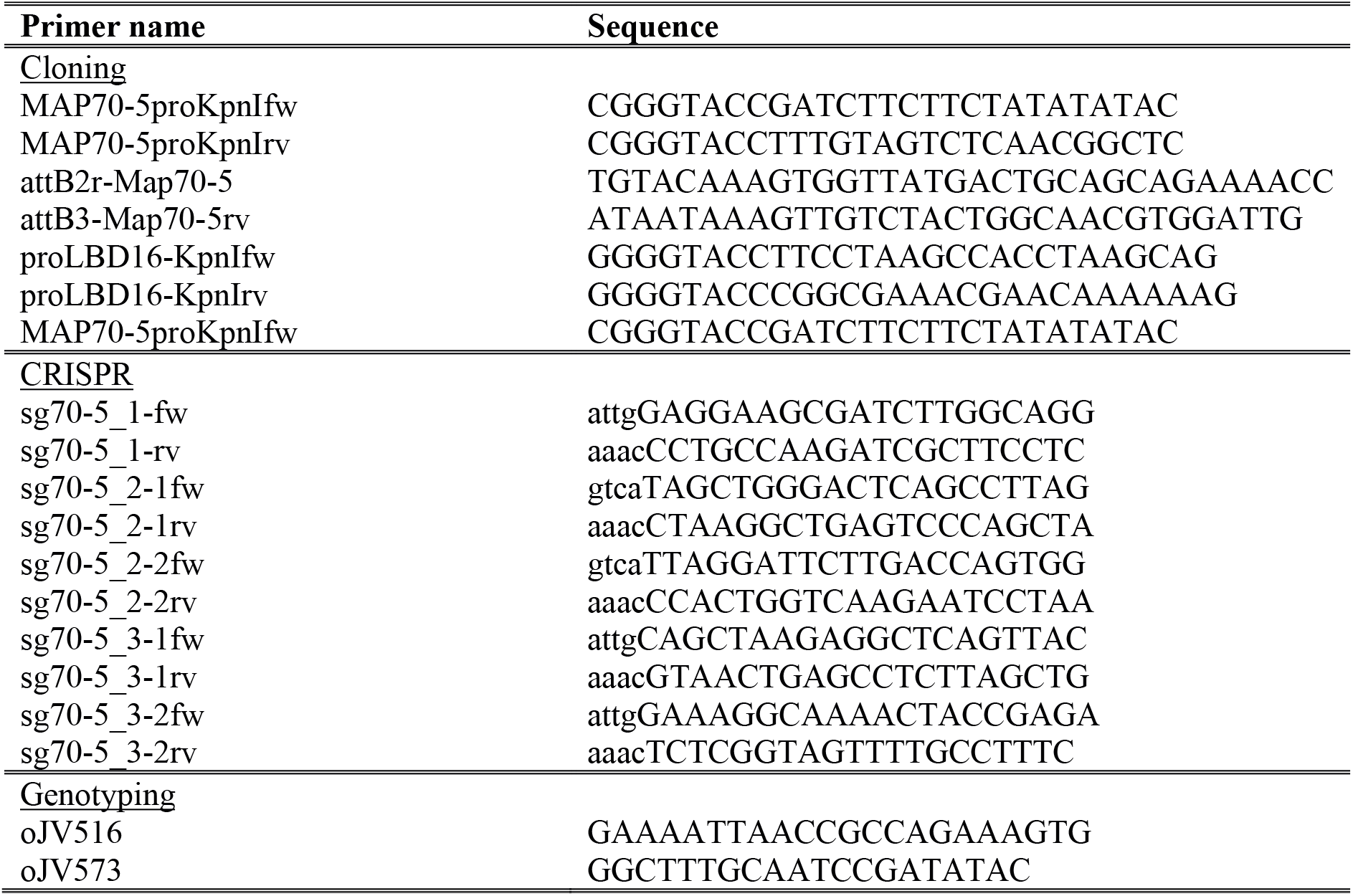
Primers used for cloning, CRISPR and genotyping.

## Notes

### Competing Interest Statement

The authors have declared no competing interest.

